# Abnormal enteric nervous system organization and gastrointestinal motility in mice with valproic acid-induced neural tube defects

**DOI:** 10.64898/2026.02.26.708353

**Authors:** Gabrielle R. Barsh, Kelbi Banducci, Julia A. Kaltschmidt

## Abstract

**Background:** Neurogenic bowel is a major cause of morbidity in patients affected by neural tube defects (NTDs) such as spina bifida, but the underlying reasons for bowel dysfunction are unknown. An absolute requirement for gastrointestinal (GI) motility is the enteric nervous system (ENS) located within the walls of the GI tract. Enteric neurons coalesce into circumferential stripes throughout embryonic and early postnatal development, and this gradual organization of the ENS into enteric neuronal stripes correlates with the emergence of neurogenic GI motility. We hypothesized that NTDs are associated with changes in ENS organization that correlate with specific GI motility defects.

**Methods:** We used prenatal valproic acid (VPA) exposure as a model for NTDs in embryonic mice. We used immunohistochemistry, high resolution confocal imaging, and *ex vivo* motility assays to assess enteric neuronal stripes and gastrointestinal motility in embryos with a VPA-induced neural tube defect.

**Key results:** GI tracts from embryos with a VPA-induced NTD contain blood. Structurally, the enteric neuronal stripes are thinner with a narrower interstripe distance, leading to an increased number of stripes. Functionally, GI motility is abnormal, with increased contraction frequency and increased length of contractile segments.

**Conclusions and inferences:** ENS organization and GI motility are disrupted in mouse embryos with a VPA-induced NTD. This has important implications for our understanding of neurogenic bowel in central nervous system diseases such as spina bifida.

**Key Points:**

- VPA exposure is a reliable model of neural tube defects with variable intralitter susceptibility
- Embryos with a VPA-induced neural tube defect have blood in the amniotic sac and within the lumen of the gastrointestinal tract
- Enteric nervous system organization is abnormal in the duodenum and jejunum of embryos with a VPA-induced neural tube defect, with thinner enteric neuronal stripes and narrower interstripe distance
- *Ex vivo* gastrointestinal motility is abnormal in the duodenum and jejunum of embryos with a VPA-induced neural tube defect, including increased contraction frequency and increased length of the contractile segment

## Introduction

Neural tube defects (NTDs) are the most common congenital malformation of the central nervous system, affecting hundreds of thousands of pregnancies annually worldwide^1^. Neurogenic bowel, broadly defined as bowel dysfunction caused by neurologic disease, is a leading cause of morbidity among patients with NTDs such as spina bifida^2,3^. However, despite the known morbidity associated with neurogenic bowel, mechanisms underlying bowel dysfunction are poorly understood.

While neurogenic bowel commonly occurs in diseases affecting the central nervous system (CNS), recent attention has been given to the enteric nervous system (ENS), the part of the peripheral nervous system that resides within the gut wall and is responsible for gastrointestinal (GI) motility^4^. In patients with spina bifida, ENS nerve fibers are thinner and enteric neuron density is decreased^5^. To address whether the ENS could be contributing to the GI dysfunction associated with NTDs, we set out to assess ENS organization and concomitantly characterize GI motility in a mouse model of NTDs.

The ENS consists of two distinct neuronal plexuses that reside within the walls of the GI tract. Neurons in the myenteric plexus are grouped into ganglia that are intercalated between two muscle layers and control GI motility^6^. We have previously shown that myenteric ganglia arrange into circumferential stripes during embryonic development, and that enteric neuronal stripe organization is distinct throughout the different regions of the GI tract in both mouse and human^7,8^. In mice, myenteric neuron birth peaks from embryonic day 10.5 (E10.5) to E14.5^6^. Neurons are initially scattered, but slowly coalesce into progressively more defined stripes. Enteric neuronal stripes become distinct in the proximal GI tract by E18.5 and in the distal GI tract during postnatal development^8^. The timing of enteric neuronal stripe formation correlates with the emergence of coordinated neurogenic GI motility in the small intestine, with clustered ripples present at E16.5 and propagating contraction complexes present by E18.5^9,10^. We therefore asked whether enteric neuronal stripes are disrupted in mice with NTDs, thereby potentially contributing to GI dysfunction.

A well-known cause of NTDs is exposure to teratogenic medications such as valproic acid (VPA), which is commonly used to treat seizures, migraines, and bipolar disorder^11^. VPA exposure during pregnancy increases the risk of NTDs 5-20 fold^12,13^. Gestational VPA exposure is also a well-established model of NTDs in rodents^14–17^. In mice, exposure to VPA during neural tube development induces NTDs with an exencephaly phenotype^14,18^. Prenatal VPA exposure in rats has been associated with altered structure of the ENS, as well as slow GI motility, although the pups do not have NTDs^19,20^.

Here, we characterize a mouse model of VPA-induced NTDs and show that in the small intestine, ENS organization and GI motility are altered. Enteric neuronal stripes in the duodenum and jejunum are increased in number with decreased interstripe distance, and increased GI contractile segment length. The duodenum, additionally, has thinner neuronal stripes and increased GI contraction frequency.

## Material and Methods

### Mice

All procedures adhered to the National Institutes of Health Guide for the Care and Use of Laboratory Animals and were approved by the Stanford University Administrative Panel on Laboratory Animal Care. Adult mice were housed up to 5 animals per cage, given food and water ad libitum, and maintained in a 12 hour/12 hour light/dark cycle. All experiments were performed on Swiss Webster mice (Charles River CFW[SW], strain code 024). For each experiment, we used a minimum of 2 control litters and 2 experimental litters per age. Experiments included both males and females, determined by genotyping for presence of Y chromosome with the following primers: 5’-GAC TAG GTT CAT AGG CAC TGG-3’ (X forward), 5’-CCG CCA AAA CTC CTT CTC TAC-3’ (X reverse), 5’AAA GGA ATT CTG GAG GCT GG-3’ (Y forward), 5’TCT TTG GAG TGA ATG TCT GCT C-3’ (Y reverse). Given the variability in litter size, no a priori sample size calculation was performed.

### Valproic acid injection and dissection

To date embryonic age, we used timed breeding and inspected females each morning for vaginal plugs. Plugged females were removed from the breeding cage. Females were weighed on the day of copulation and again at E8.5, and if they gained >2.5g of weight they were deemed likely pregnant and suitable for injection. Pregnant dams received an intraperitoneal injection at E8.5 of either sterile water (control) or 400mg/kg of valproic acid sodium salt (Sigma-Aldrich) dissolved in sterile water. The timing and dose of VPA were selected based on prior literature showing reliable induction of exencephaly^17,21,22^.

Embryos were collected at E16.5 or E18.5, as specified. Pregnant mice were culled by CO_2_ and cervical dislocation. The uterus was removed and each implantation site was inspected visually to determine number and location of implantations for each litter, including uterine horn (left versus right) and uterine position (ovarian versus middle versus cervical) as previously described^17^. Each embryo was inspected visually for presence of blood in the amniotic sac. Embryos were removed from the amniotic sac, inspected visually for presence of exencephaly, and placed in a Sylgard 170 plate containing either phosphate buffered saline (PBS) on ice (for immunohistochemistry experiments) or warm carbogenated Krebs solution (for *ex vivo* motility monitor experiments). The tail of each embryo was collected to obtain DNA for genotyping. Intestines were dissected from the embryo and mesentery was carefully removed. For motility monitor experiments, a maximum of eight embryos per litter was utilized to ensure intestines could be quickly transferred to a warm organ bath with circulating carbogenated Krebs solution. For all other experiments, intestines were pinned to the Sylgard dish using any remaining mesentery and pulled taught for fixation. Tissue was fixed in 4% paraformaldehyde for 90 minutes at 4°C. The length of the cleaned intestine was measured with an adjacent ruler.

### Immunohistochemistry and image acquisition

For whole mount imaging, fixed intestines were placed into PBT (PBS, 1% bovine serum albumin, 0.1% Triton X-100) containing primary antibodies overnight at 4°C with shaking. The following day, intestines were washed three times in PBT at room temperature (RT) for 5 minutes each, then placed in PBT containing secondary antibodies for two hours at RT with shaking. Intestines were washed twice more in PBT for 5 minutes each and transferred to PBS at RT. ∼1cm segments from each region of the intestine were selected for mounting. Duodenum was defined as the most proximal part of the small intestine, jejunum was the middle section of the small intestine, and ileum was the most distal part of the small intestine. Colon was divided into proximal and distal segments. Each section of the intestine was cannulated with a cleaning wire for 33-gauge needles (Hamilton) and cut open along the mesentery. Intestines were mounted as full-thickness preparations onto slides with Fluoromount-G (Southern Biotech).

For cross section imaging, fixed intestines were placed in 30% sucrose overnight at 4°C. 500mm sections of each region (duodenum, jejunum, ileum, proximal colon, distal colon) were cut out, embedded in OCT as a tube, and stored at -80°C. Tissue sections were cut at 14μm thickness with a Leica cryostat CM3050 S and placed on slides. Tissue was then rehydrated in PBS and stained as above.

Primary antibodies used were rabbit anti-HuC/D (1:2000, Abcam 184267), rat anti-E-cadherin (1:1000, ThermoFisher 13-1900), rat anti-CD31 (1:500, BD Pharmingen 557355), and goat anti-smooth muscle actin (1:2000, Abcam 21027). Fluorophore-conjugated secondary antibodies were used at 1:500 to visualize signal (Jackson Labs or Invitrogen).

Images were acquired using a 20x oil objective on a Leica SP8 confocal microscope. Tiled images were identified, acquired, and stitched using the Navigator mode within LASX (Leica). A z stack of the entire region was taken, ensuring full thickness was captured in all tiles. Z stacks were acquired with 2μm between each focal plane.

### H&E stain and image acquisition

Intestines were dissected and fixed in the same manner as for immunohistochemistry experiments. After fixation, intestines were placed in 30% sucrose overnight at 4°C. Each region was mounted in OCT, frozen, and sectioned with a Leica cryostat as above. Slides were stained with hematoxylin and eosin (H&E), cleared with xylene, and mounted with DPX mountant as previously described^23^. Brightfield images were taken on a Zeiss Axioskop 2 plus.

### Image analysis

Image analysis was performed using ImageJ/FIJI (NIH, Bethesda, MD). All image analysis was done blinded to condition. Assays for number of neurons, interstripe distance, and number of stripes were performed on 500x2000μm maximal intensity projections. To determine number of neurons, we adapted previously published COUNTEN software to output total number of neurons, total number of ganglia, and neuron count per ganglia^24^. To determine number of stripes and interstripe distance, images were blurred and thresholded in FIJI, and a grid of 100x100μm regions was projected onto the image using the grid tool in FIJI. Along each horizontal line of the grid (5 lines total per image), the line segment tool was used to mark the midpoint of each neuronal stripe intersected. A custom FIJI macro was used to output number of stripes and interstripe distances along each line. Interstripe distances were averaged along each line, and those values were averaged to determine a single value for each embryo. For number of stripes, we used the median number along each line, and then calculated the median of those values to determine a single value for each embryo.

To quantitatively assess ENS organization, we used a previously published code for assessing enteric neuronal cytoarchitecture^8^. We analyzed 4 nonoverlapping 400x400μm tiles per sample in duodenum and jejunum, and up to 4 nonoverlapping tiles per sample in ileum, proximal colon, and distal colon. Fewer tiles were used in ileum, proximal colon, and distal colon due to the smaller size of those regions compared to duodenum and jejunum. Tiles served as technical replicates, with values averaged to determine a single value for each embryo. To assess neuron distribution, we used nearest-neighbor distance measurements to determine how the measured distribution differed from a hypothesis of random neuron positions, i.e. complete spatial randomness, as previously described^8^. To determine stripe width, we used conditional intensity function analysis to generate a spatial density map of neuron location, which was then transformed into a one dimensional line to determine stripe width as previously described^8^.

### *Ex vivo* motility analysis

We used the *ex vivo* motility monitor as previously described^9^. Briefly, intestines were submerged in an organ bath containing circulating carbogenated 32°C Krebs solution (pH 7.4 containing (in mmol L^-1^): 117 NaCl, 4.7 KCl, 3.2 CaCl_2_ (2H_2_O), 1.5 MgCl_2_ (6H_2_O), 25 NaHCO_3_, 1.2 NaH_2_PO_4_ and 11 Glucose) and a heated water bath below to maintain chamber temperature. The intestines were pinned taut on Sylgard 170 in the chamber base. Visualization of intestine from embryos with a VPA-induced NTD was limited, due to the reduced contrast between the blood in the intestinal lumen and the black Sylgard background. We therefore added a flat polystyrene plastic sheet to the organ bath to improve contrast from background, using white for intestines that contained blood and black for intestines that did not contain blood. Intestines were allowed to acclimate in the organ bath for 10 minutes undisturbed. 10-minute videos were recorded at 8 frames/second using FlyCap2 capture software (Imaging Source) and a high-resolution monochromatic firewire industrial camera (Imaging Source, DMK41AF02) connected to a 2/3′′ 16mmf/1.4 C-Mount Fixed 629 Focal Lens (Fujinon HF16SA1) mounted above the organ bath.

Spatiotemporal maps (STMs) were generated using Scribble 2.0 and Matlab (2014b) plugin Analyze 2.0^25,26^. STMs were cropped to analyze 1cm of intestine length in either the duodenum or the jejunum and included the entire 10-minute video. Contractions were identified using the ridge detection filter in FIJI to determine number and length of contractile segments across the entire STM. Contractile segment length was averaged to determine a single value for each STM. To determine contraction frequency, we used a custom FIJI macro to bin the STM into 10 regions of 1mm each and assessed number of contractions in that region divided by time length of video. The frequency in each of the 10 bins was averaged to determine a single value for contraction frequency for each STM.

### Statistical analysis

Statistical tests and graphical representation of data were done using Prism 10 software (GraphPad). Statistical comparisons for categorical data (NTD outcome in response to VPA exposure, presence of blood in the GI tract, sex genotype, left versus right uterine horn) were performed using chi-square analysis. Statistical comparison for uterine position was done using Fisher’s exact test given the smaller sample size. Statistical comparisons for all other numerical data were performed using either Welch’s t test (for two group comparisons) or one-way ANOVA (for multiple group comparisons) followed by Tukey’s multiple comparisons test to further investigate differences between individual groups.

## Results

### Exposure to valproic acid causes NTDs with variable intralitter susceptibility

To assess the impact of NTDs on ENS organization and GI motility, we adapted a previously published mouse model of VPA exposure in Swiss Webster mice during gestation^17,21,22^. We found that administering a single dose of 400mg/kg VPA on embryonic day 8.5 (E8.5) led to exencephaly in about one third of the exposed embryos when assessed at either E16.5 (Fig. 1D) or E18.5 (Fig. 1A, E), similar to previously published work^21^. While the percentage of affected embryos varied considerably in individual litters, nearly all litters had at least one affected embryo (Fig. 1 F, G). There were also a significant number of post-implantation losses (Fig. 1 D-G), consistent with prior findings that VPA exposure leads to gestational and perinatal lethality^17,27^.

**Figure 1.**
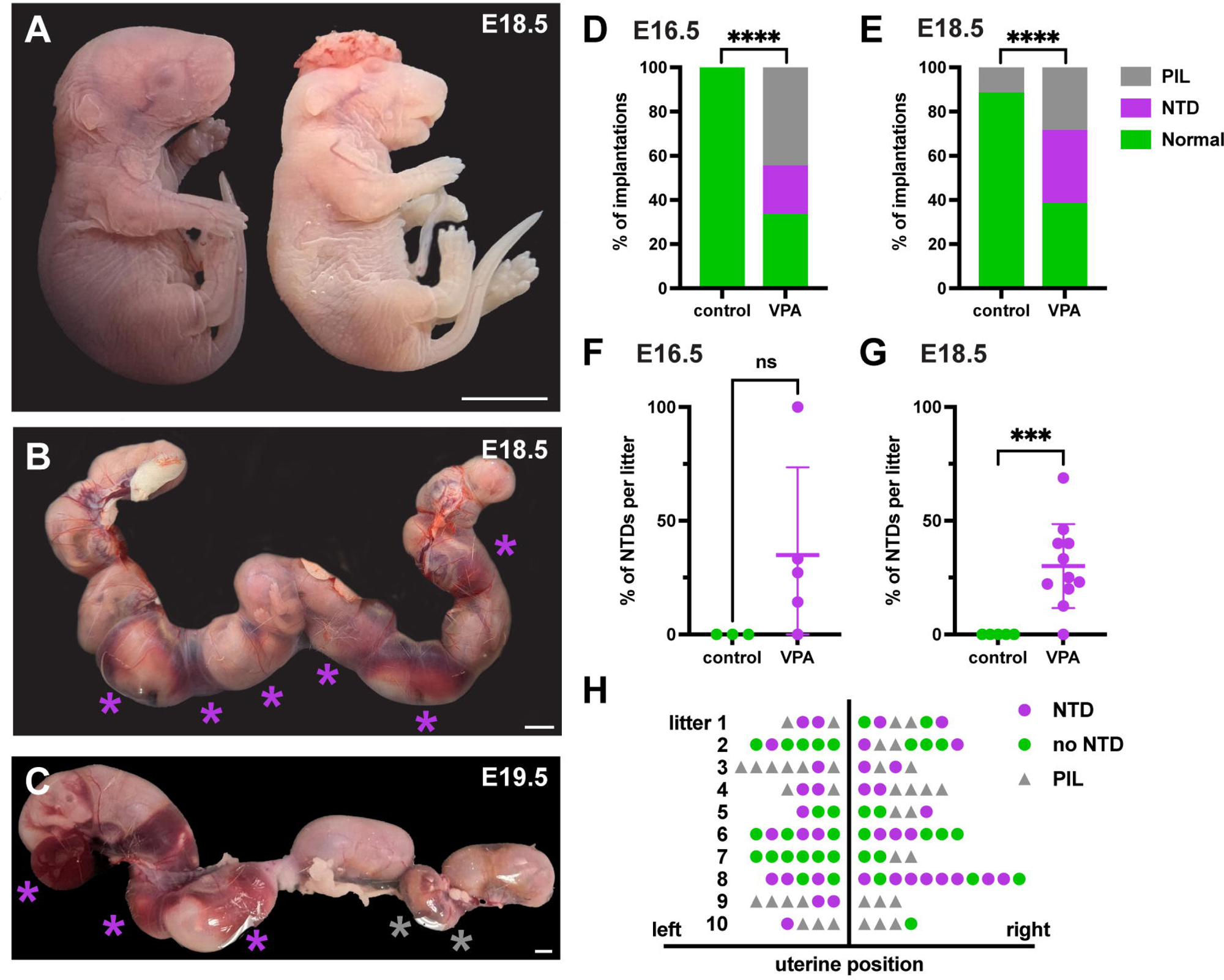
Outcomes of embryos exposed to valproic acid. (A) Two embryos at E18.5 from a litter exposed to valproic acid (VPA). Left, unaffected embryo. Right, embryo with exencephaly, a type of NTD. (B, C) Uterus containing a litter exposed to VPA dissected at E18.5 (B) or E19.5 (C). Purple asterisks denote embryos affected by an NTD, gray asterisks denote post-implantation losses. Amniotic sacs of affected embryos contain blood. (D, E) Exencephaly outcomes of embryos at E16.5 (D) or E18.5 (E). 22% of embryos have an NTD at E16.5, 33% have an NTD at E18.5. D, control N = 2 litters and n = 26 implantations; VPA N = 5 litters and n = 42 implantations, p<0.0001. E, control N = 6 litters and n = 68 implantations; VPA N = 11 litters and n = 111 implantations, p<0.0001. Data displayed as percentage of total implantations, statistics performed on raw total number of implantations per condition. (F, G) Outcomes of embryos exposed to VPA shown by percentage of litter affected at E16.5 (F, VPA mean 34.97%) or E18.5 (G, VPA mean 30.10%). (H) Schematic showing uterine position of affected embryos and post-implantation losses at E18.5. VPA, valproic acid. NTD, neural tube defect. PIL, post-implantation loss. Scale bars, 0.5cm. ***p < 0.001, ****p<0.0001.

We next asked whether the observed variable intralitter susceptibility to VPA teratogenicity is due to embryo sex or intrauterine position. We found that females were more likely to have exencephaly compared to males (58.33% of females affected compared to 33.33% of males, p = 0.04, Fig. S1A-C). Neither intrauterine position nor left/right uterine horn location affected likelihood of NTD (Fig. 1H, Fig. S1D-G). This agrees with previously published work in mice that found female embryos are more susceptible to VPA compared to males^17^. Overall, these results confirm our usage of 400mg/kg VPA prenatal exposure as a reliable mouse model of NTDs.

### Embryos with a VPA-induced NTD have blood inside the gastrointestinal tract

We next asked whether GI gross morphology is affected in embryos with a VPA-induced NTD (referred to as VPA-NTD). We compared VPA-NTD embryos and embryos from a control dam that was injected with sterile water (referred to as controls). We additionally considered embryos exposed to VPA without an NTD (unaffected littermates, referred to as VPA-noNTD), reasoning that this group would allow us to determine whether any observed differences were related to presence of an NTD specifically, rather than VPA exposure.

We first assessed GI length and noticed variability in the length of the small intestine in embryos exposed to VPA. At E16.5, the small intestine from VPA-NTD embryos is shorter in length compared to VPA-noNTD embryos, but not significantly different from control embryos (Fig. 2C). This difference disappears at E18.5, and instead intestines from VPA-NTD embryos are significantly longer than intestines from VPA-noNTD embryos, and no different from control embryos (Fig. 2E). There is no difference in colon length at either E16.5 or E18.5 across conditions (Fig. 2D, F). Overall, there is no significant difference in the total length of the GI tract in VPA-NTD embryos compared to control embryos.

**Figure 2.**
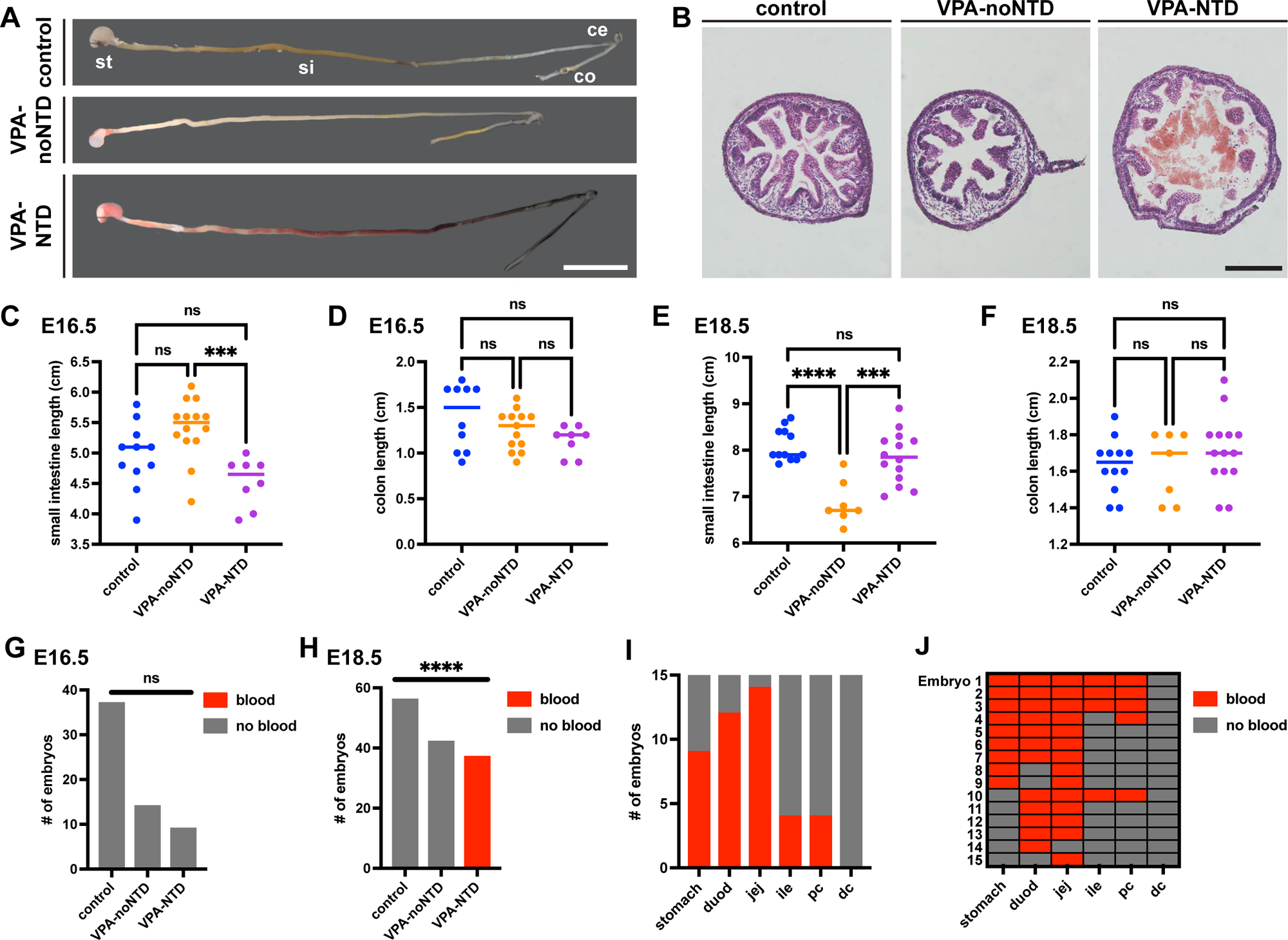
Gross morphology of the gastrointestinal tract in embryos exposed to valproic acid. (A) Gastrointestinal (GI) tracts at E18.5 from control (top), VPA-noNTD (middle), or VPA-NTD embryo (bottom). (B) Sections of ileum from control (left), VPA-noNTD (middle), or VPA-NTD embryo (right) at E18.5. Note presence of blood within the lumen in image on the right. (C, D) Quantification of small intestine (C) and colon (D) length at E16.5. (E, F) Quantification of small intestine (E) and colon (F) length at E18.5. (G, H) Quantification of blood within the lumen of the GI tract at E16.5 (G) and E18.5 (H). (I) Distribution of intraluminal blood in VPA-NTD embryos at E18.5. (J) Presence of blood in regions of the GI tract for individual VPA-NTD embryos at E18.5. Some embryos, such as embryo 1, have blood present throughout contiguous regions. Others, such as embryo 8, have blood present in discontiguous regions. St, stomach; si, small intestine; ce, cecum; co, colon; duod, duodenum; jej, jejunum; ile, ileum; pc, proximal colon; dc, distal colon. Scale bars, 1cm (A), 100μm (F). ***p < 0.001, ****p<0.0001.

While assessing the GI tract of VPA-NTD embryos, we noted blood within the GI lumen (Fig. 2A, B). We found intraluminal blood in every VPA-NTD embryo at E18.5 (Fig. 2H). In comparison, we did not detect blood in the GI tract of VPA-NTD embryos at E16.5 or VPA-noNTD embryos at either age (Fig. 2G, H). Within the GI tract of VPA-NTD embryos at E18.5, blood was predominantly found in the proximal regions, including the stomach (Fig. 2I). While the majority of embryos (80%) had blood in contiguous segments throughout the length of the GI tract, some had discontiguous areas of blood (Fig. 2J).

To assess the origin of the intraluminal blood in VPA-NTD embryos, we explored two possibilities: intraluminal intestinal bleeding versus swallowed blood. We first performed immunohistochemistry on cross-sectional images to assess whether the layered structure of the GI tract appeared typical and blood vessels were present. We did not find any major differences in the gross structure of the GI tract between control, VPA-noNTD, and VPA-NTD embryos, with a corrugated epithelium (Fig. S2B) and blood vessels present within villi (Fig. S2A) in all conditions.

We next assessed the amniotic sacs as an alternative blood source, and found blood in the amniotic sacs of VPA-NTD embryos, but not control or VPA-noNTD embryos (Fig. 1B, C). Amniotic hemorrhage was present as early as E16.5, preceding intraluminal GI blood by two days. Given the presence of amniotic hemorrhage, together with observations from the literature that mouth opening occurs in mouse embryos as early as E14.5^28^, we posit embryos may be swallowing blood from the amniotic sac.

### Enteric neuron organization is disrupted in embryos with a VPA-induced NTD

We next asked whether ENS structure is disrupted in VPA-NTD embryos. Enteric neuronal stripes become progressively more established throughout embryonic and postnatal development^8^. We used immunohistochemical labeling of HuC/D to visualize enteric neuronal stripes at E18.5^8,29^. In control embryos, stripes are relatively well-established in the duodenum and jejunum (Fig. 3A, B, and reference 8) while they are less well-formed in the other regions of the GI tract (Fig. S3A-C, and reference 8). In VPA-NTD embryos, as well as VPA-noNTD embryos, enteric neuronal stripes are visually present in the duodenum and jejunum (Fig. 3A, B).

**Figure 3.**
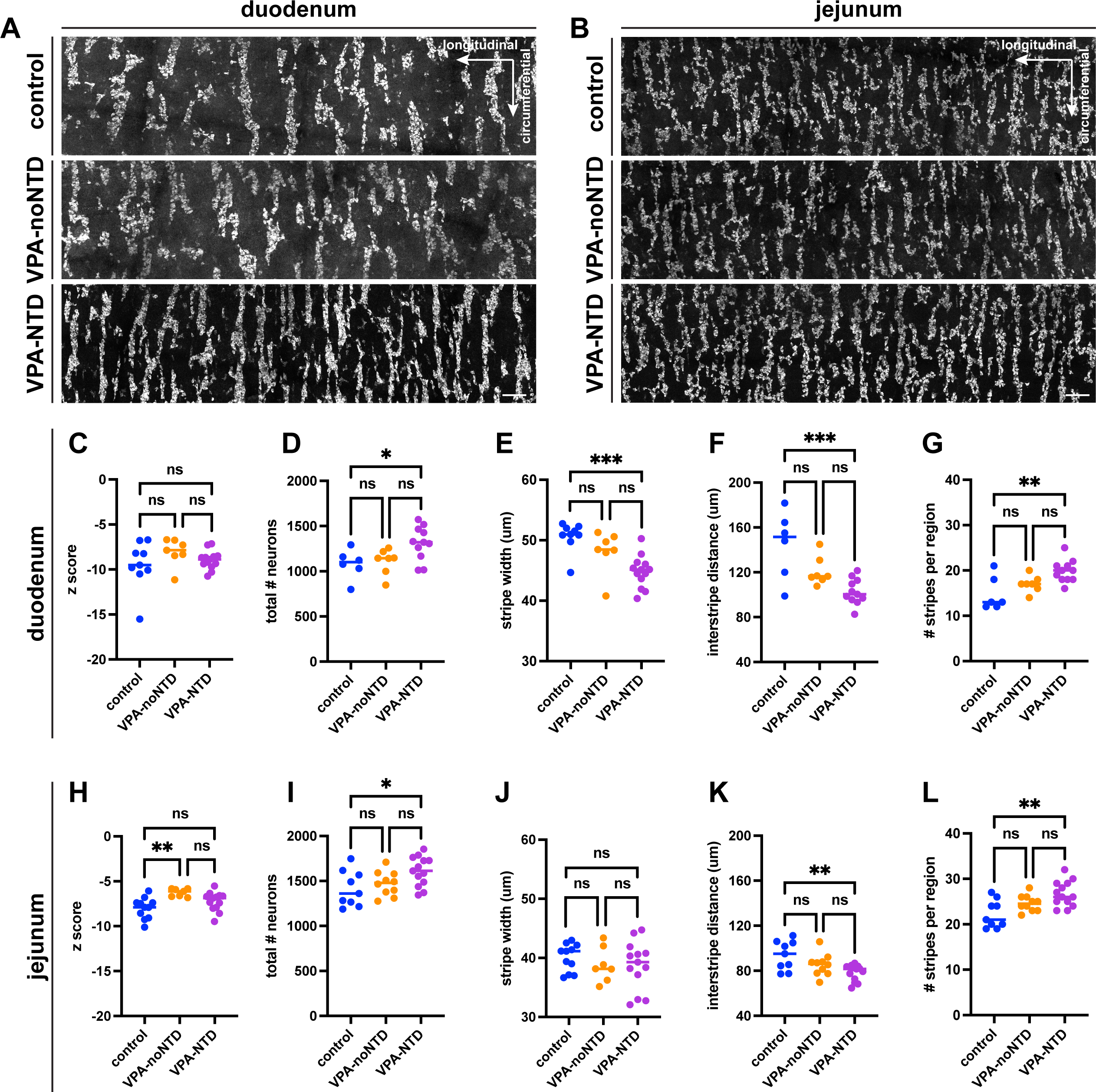
Enteric neuronal stripes in proximal intestine in embryos exposed to valproic acid. (A, B) Sections of duodenum (A) or jejunum (B) in a whole mount preparation showing enteric neurons labeled with HuC/D in control (top), VPA-noNTD embryos (middle), or VPA-NTD embryos (bottom). (C-L) Quantitative assessments of neuronal stripe organization in duodenum (C-G) or jejunum (H-L). (C, H) Assessment of neuron distribution in either duodenum (C) or jejunum (H). Negative z score indicates more clustering than random. Positive z score indicates less clustering than random. (D, I) Quantification of neuron density in the duodenum (D) and jejunum (I). (E, J) Quantification of stripe width in the duodenum (E) and jejunum (J). (F, K). Quantification of interstripe distance in duodenum (F) and jejunum (K). (G, L). Quantification of stripe density in the duodenum (G) and jejunum (L). *p<0.05, ** p<0.01, ***p<0.001. Scale bar, 100μm.

To quantitatively assess neuronal distribution within enteric neuronal stripes, we used nearest-neighbor distance measurements^8^. In VPA-NTD embryos, neurons in both the duodenum and jejunum are highly clustered consistent with striped patterning, similar to control embryos (Fig. 3C, H). In VPA-noNTD embryos, neurons in the duodenum are also highly clustered (Fig. 3C), while neurons in the jejunum are slightly more dispersed compared to control embryos (Fig. 3H). We also assessed more distal parts of the intestine where enteric neuronal stripes are still being established. In the ileum, proximal colon, and distal colon, neurons remain relatively dispersed with no significant difference across conditions (Fig. S3D, F, G). Together these results provide quantitative evidence that enteric neuronal stripes are present in the duodenum and jejunum of VPA-NTD embryos.

We next asked whether there was a difference in the detailed organization of the enteric neuronal stripes. We first used COUNTEN software^24^ to assess the overall number of enteric neurons. In VPA-NTD embryos, the number of neurons is ∼20% increased in both the duodenum and jejunum as compared to VPA-noNTD and control embryos (Fig. 3D, I). There is no difference in neuron number across groups in more distal tissues including ileum or distal colon (Fig. S3 E, H).

We next assessed enteric neuronal stripe width, interstripe distance, and number of stripes in VPA-NTD embryos. To assess stripe width, we used conditional intensity function analysis to create a spatial probability map of neuron location^8^. We found that stripes are narrower in the duodenum of VPA-NTD embryos as compared to controls (VPA-NTD mean 44.8μm versus control mean 50.6μm, Fig. 3E), though this difference is not seen in the jejunum (Fig. 3J). The interstripe distance is ∼30% shorter in the duodenum and ∼20% shorter in the jejunum of VPA-NTD embryos as compared to control embryos (Fig. 3F, K). This results in an increased number of stripes within a given area in VPA-NTD embryos in both the duodenum (∼30% increase, Fig. 3G) and the jejunum (∼20% increase, Fig. 3L). There is no difference in stripe width, interstripe distance, or number of stripes in VPA-noNTD embryos compared to control embryos. Together, these results show that enteric neuronal stripe organization is altered in embryos with a VPA-induced NTD.

### Gastrointestinal motility is altered in embryos with a VPA-induced NTD

Given that ENS structure is disrupted in VPA-NTD embryos, we next asked whether GI function is also affected. We utilized an *ex vivo* embryonic motility monitor, in which the intestines are dissected and placed into a warm buffered bath to visualize contractions^9,10^ (Fig. 4A, B). In this setup, central nervous system input to the intestine is severed, such that any contractions seen can be attributed to autonomous ENS activity. We have previously used this *ex vivo* setup to quantitatively assess neurogenic motility in embryonic intestines through generating spatiotemporal maps (STMs) of intestinal movement^9^. When the intestine contracts, the diameter narrows, seen as a horizontal black line on the STM (Fig. 4C, D). At E18.5, in intestines from control embryos, we found that contraction frequency in the duodenum is about 2 contractions per minute, similar to previously published values^9,10^ (Fig. 4E). However, in VPA-NTD embryos, contraction frequency in the duodenum is increased about two-fold compared to control embryos (Fig. 4E, mean 2.5 contractions/minute in control embryos versus 4.6 contractions/minute in VPA-NTD embryos). There was no difference in contraction frequency in the jejunum of VPA-NTD embryos (Fig. 4G). We next measured the average length of the contractile segments and found that in VPA-NTD embryos it is ∼50% longer in the duodenum (Fig. 4F, mean 0.44cm in controls versus 0.68cm in VPA-NTD embryos) and ∼80% longer in the jejunum (Fig. 4H, mean 0.40cm in controls versus 0.74cm in VPA-NTD embryos), as compared to control embryos. We saw no GI motility differences in VPA-noNTD embryos as compared to controls. Together, these results show that embryos with a VPA-induced NTD have altered GI motility, with increased contraction frequency and increased length of contractile segments.

**Figure 4.**
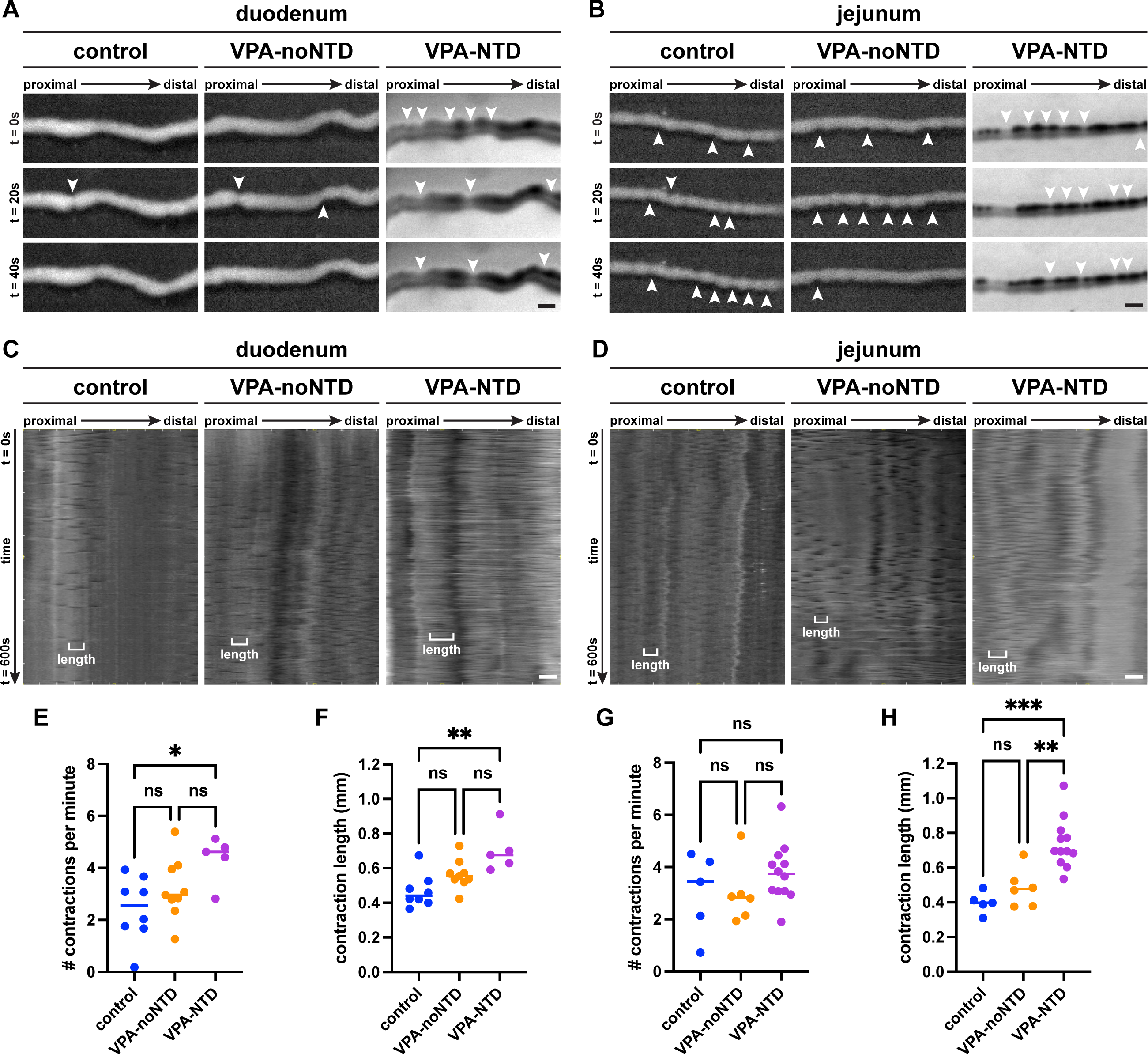
Neurogenic motility in embryos exposed to valproic acid. (A, B) Representative timelapse images of the duodenum (A) or jejunum (B) in an *ex vivo* motility monitor showing contraction patterns in control embryos (left), VPA-noNTD embryos, (middle), or VPA-NTD embryos (right). Arrowheads denote contractions. (C, D) Representative spatiotemporal maps of GI motility in the duodenum (C) or jejunum (D) across conditions. Maps are in grayscale with darker gray indicating narrower diameter and lighter gray indicating wide diameter, such that contractions appear as horizontal black lines. An example of a single contraction is shown on each map denoting length of contractile segment. (E, F) Quantification of contraction frequency (E) and length of contractile segment (F) in the duodenum. (G, H) Quantification of contraction frequency (G) and length of contractile segment (H) in the jejunum. *p<0.05, **p<0.01, ***p<0.001. Scale bar, 1mm.

## Discussion

Neurogenic bowel is a phenotype that occurs in many diseases of the central nervous system, including NTDs such as spina bifida. Here, we show the organization of the ENS is affected in a mouse model of NTDs. We characterize concomitant GI motility defects, and suggest that the ENS might play a direct role in neurogenic bowel.

We used gestational exposure to VPA as a model of NTDs, which is a clinically salient model given the prevalence of women exposed to VPA before and during pregnancy^30^. Although VPA exposure is a well-established model of NTDs, it is only recently that variable intralitter susceptibility to VPA has been described. Brown and colleagues found that females were more susceptible to VPA-induced NTDs at 600mg/kg, but not at the lower dose of 400mg/kg^17^. In our work, we find that females are more susceptible at the relatively lower dose of 400mg/kg. This difference may be attributed to the use of Swiss Webster (our work) versus CD-1 mice (reference 17), as different strains vary in their susceptibility to VPA teratogenicity^14,31^. In human patients, NTDs are more prevalent in females than males^32,33^, although it is unknown whether VPA specifically is more teratogenic in females.

We centered our analysis of ENS organization on enteric neuronal stripes. We previously showed that the emergence of enteric neuronal stripes is correlated with the development of coordinated neurogenic GI motility^9^. Our results here show altered structure of enteric neuronal stripes and concomitantly altered GI motility. We focused much of our analysis on the duodenum and jejunum, as these are the regions in which enteric neuronal stripes are present during normal development at the time point analyzed in this study. While we find changes in ENS organization and GI motility in VPA-NTD embryos in both the duodenum and jejunum, effects vary between these two regions. Enteric neuronal stripes are narrower in the duodenum of VPA-NTD embryos, but not in the jejunum. In control embryos, enteric neuronal stripes in the jejunum are narrower compared to the duodenum, so it is possible that stripes in the jejunum are too narrow for us to detect subtle differences in stripe width in VPA-NTD embryos. Similarly, we detect an increase in contraction frequency in the duodenum but not in the jejunum. This raises the intriguing possibility that specific features of enteric neuronal stripe organization are important for specific aspects of motility patterns—for example, that stripe width influences contraction frequency, given those two parameters are affected in the duodenum of VPA-NTD embryos. It remains to be determined whether other factors such as changes in cell type distribution or neuronal connectivity are also altered in VPA-NTD embryos and thereby could contribute to altered motility patterns.

The differences that we see in neuron number, enteric neuronal stripe structure, and GI motility are specific to embryos with an NTD, and not seen in embryos exposed to VPA without an NTD. This is an important distinction, as it is well known that VPA can influence multiple different aspects of embryonic development beyond neural tube development, such as craniofacial structures and cardiac development^12,21^. This raises the potential criticism that the differences we see in VPA-NTD embryos could be a result of tissue-wide VPA exposure. However, in the majority of our analyses of ENS structure and GI function, there is no difference in VPA-noNTD embryos compared to control embryos, suggesting that VPA exposure alone may affect gross morphology of the GI tract but is relatively insufficient to alter ENS organization. The exception was a slightly more dispersed neuron distribution in the jejunum of VPA-noNTD embryos. It remains unclear why neurons are more dispersed in VPA-noNTD jejunums. Overall, we conclude that ENS organization and GI function are highly correlated with presence of an NTD.

How do NTDs impact ENS development? In our analyses, we found an increased number of neurons present in the duodenum and jejunum in VPA-NTD embryos. As enteric neural crest cells delaminate from the neural tube prior to migrating to the intestine, an open neural tube could theoretically impact enteric neural crest cells early in development, potentially leading to increased neurogenesis or changes in cell migration. It is important to note that GI motility is also regulated by extrinsic innervation from vagal and spinal nerves^34^, and it remains unknown whether extrinsic innervation of the GI tract is disrupted by NTDs. Future studies examining the impact of NTDs on early enteric neural crest cell development and migration or vagal and spinal innervation of the GI tract should be performed to elucidate how these processes might be affected.

An unexpected finding of our work was the presence of blood within the lumen of the GI tract in VPA-NTD embryos. Though the processes leading to intraluminal blood remain unknown, we suggest this may represent swallowed blood from amniotic fluid. The presence of amniotic hemorrhage has not been previously reported in the use of prenatal VPA exposure in mouse models, though some authors have described subcutaneous hemorrhage^21^. Although all VPA-NTD embryos had amniotic hemorrhage at E16.5, the intraluminal blood was not seen until later at E18.5. As mouth opening occurs in mouse embryos as early as E14.5^28^, we postulate that blood first appears in the amniotic sac, and is then swallowed into the GI tract. One outstanding question is whether the presence of blood within the GI tract is contributing to the phenotypes we describe in VPA-NTD embryos, particularly the increased frequency and length of GI contractions. Sensory neurons are present by E12.5^6^, and so it could be possible that enteric neurons are sensing and responding to differences in consistency and makeup between blood and meconium. In the adult guinea pig colon, mechanical properties of the intraluminal contents affect GI motility^35^. Future experiments intentionally altering components of the amniotic fluid could shed further light on how intraluminal contents affect embryonic motility.

In conclusion, we show here that ENS structure and GI function are disrupted in mice with a VPA-induced neural tube defect. This has important implications for considering the fundamental basis of neurogenic bowel in patients with central nervous system diseases, including spina bifida.

## Acknowledgements, funding, disclosures

We thank members of the Kaltschmidt lab for their valuable feedback. In particular, we would like to thank Eric Zhao for assistance implementing COUNTEN and Jacqueline Bendrick for helpful comments on the manuscript. We thank Anca Pasca and Courtney Wusthoff for helpful comments on project design. We thank the Brunet lab, especially Sasha Tsenter, for lending use of the brightfield microscope. This work was funded by NINDS Award Number K12NS098482 (GRB), NIH URISE scholarship (KB), a Stanford Maternal and Child Health Research Institute (JAK), the Wu Tsai Neurosciences Institute (JAK), the Stanford University Department of Neurosurgery (JAK), and a research grant from The Firmenich Foundation (JAK). The authors declare no relevant conflicts of interest exist.

## Figure Legends

**Figure S1.**
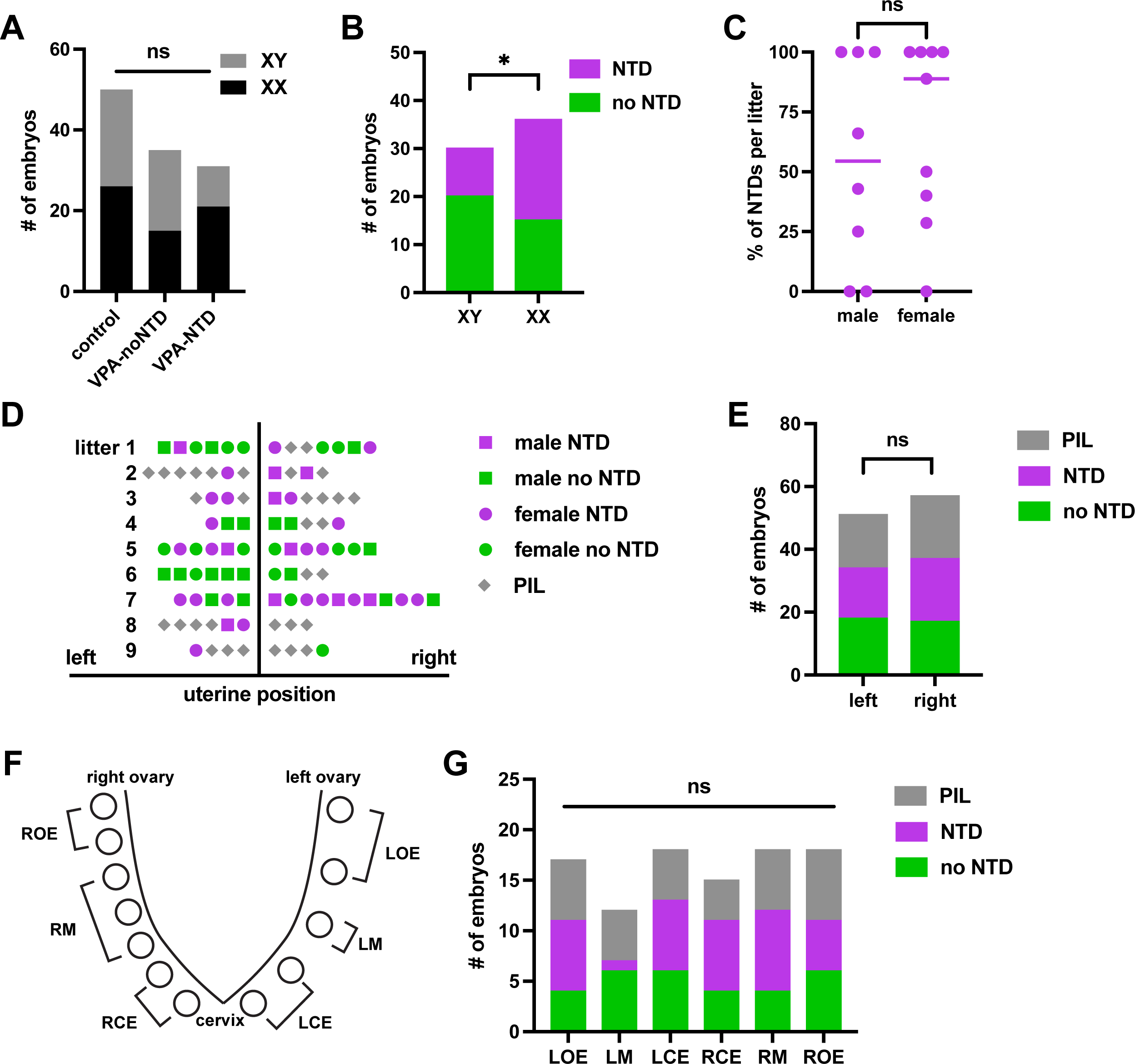
Testing factors affecting intralitter susceptibility to valproic acid-induced neural tube defects. (A) Sex genotype of embryos across experiments. Control embryos are evenly divided into males and females (52% female, 48% male), while amongst embryos exposed to VPA, those with an NTD were 68% female, 32% male, and those without an NTD were 43% female, 57% male. (B) Among VPA-exposed embryos, 33% of XY embryos have an NTD, while 58% of XX embryos have an NTD. (C) Outcomes of embryos shown by percent of litter affected. XY mean 54.2% affected, XX mean 67.5% affected. (D) Schematic showing uterine position of each affected embryo and each post-implantation loss amongst litters, with sex of each embryo denoted. (E) Outcomes of VPA-exposed embryos shown by uterine horn. (F) Schematic denoting labeling system for uterine position, inspired by reference 17. (G) Outcomes of VPA-exposed embryos shown by uterine position. VPA, valproic acid. All experiments shown in this figure were done at E18.5. NTD, neural tube defect. PIL, post-implantation loss. ROE, right ovarian end. RM, right middle, RCE, right cervical end. LCE, left cervical end. LM, left middle. LOE, left ovarian end. *p < 0.05.

**Figure S2.**
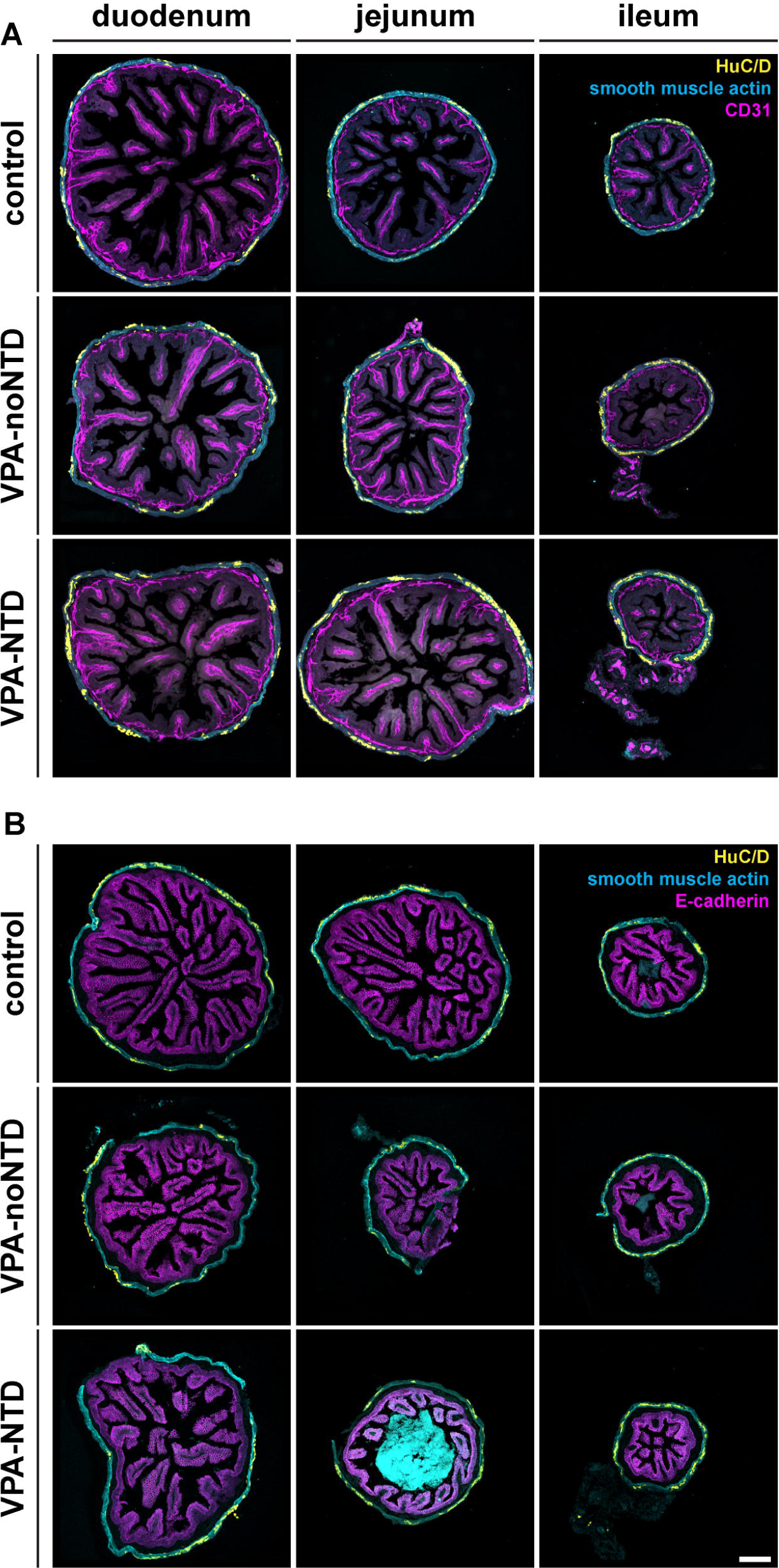
Structure of gastrointestinal tract wall in embryos exposed to valproic acid. (A, B) Cross sections of regions of the GI tract from proximal (left) to distal (right) from control (top), VPA-noNTD (middle), or VPA-NTD embryos (bottom), with immunohistochemical labeling against pan-neuronal marker HuC/D (yellow), smooth muscle actin (cyan), and either endothelial marker (CD31, magenta in A) or epithelial marker (E-cadherin, magenta in B). Scale bar, 100μm.

**Figure S3.**
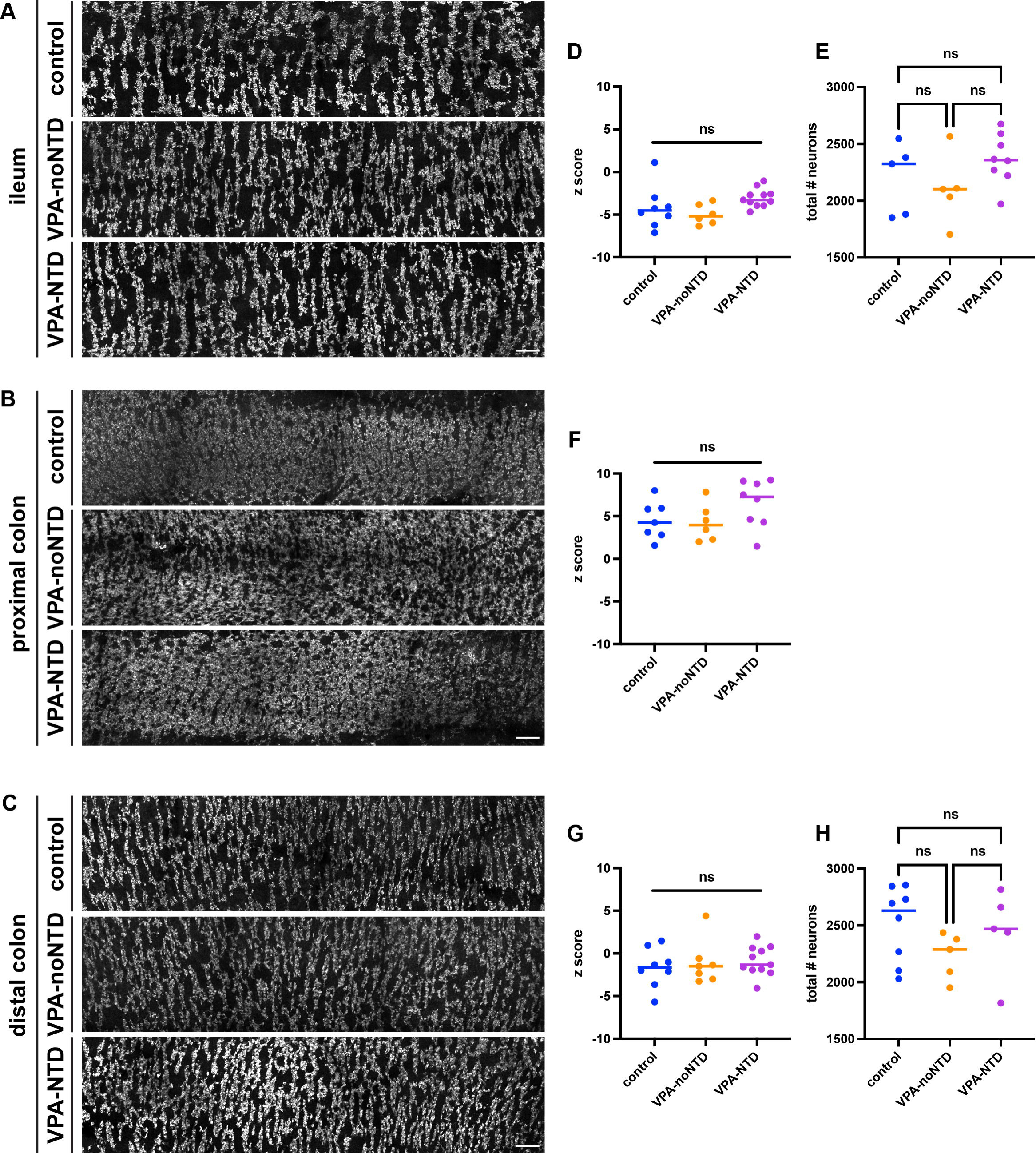
Enteric neuronal stripes in distal intestine in embryos exposed to valproic acid. (A-C) Sections of ileum (A), proximal colon (B), or distal colon (C) showing whole mount preparation with enteric neurons labeled by HuC/D in control (top), VPA-noNTD embryos (middle), or VPA-NTD embryos (bottom). (D, F, G) Z score values reflecting neuron distribution compared to spatial randomness for ileum (D), proximal colon (F), or distal colon (G). (E, H) Quantification of neuron density for ileum (E) or distal colon (H). Proximal colon omitted as the narrow diameter of the tissue limited our ability to obtain a sample large enough for analysis. Scale bar, 100μm.

